# Transplantation of GABAergic Interneuron Progenitors Restores Cortical Circuit Function in an Alzheimer’s Disease Mouse Model

**DOI:** 10.1101/2025.05.31.656412

**Authors:** Shinya Yokomizo, Megi Maci, April M. Stafford, Morgan R. Miller, Stephen James Perle, Shusaku Takahashi, Heather Brown-Harding, Linda Liang, Alex Lovely, Moustafa Algamal, Rebecca L. Gillani, Theodore J. Zwang, Douglas Richardson, Janice R. Naegele, Daniel Vogt, Ksenia V. Kastanenka

## Abstract

In addition to dementia, Alzheimer’s patients suffer from sleep impairments and aberrations in sleep-dependent brain rhythms. Deficits in inhibitory GABAergic interneuron function disrupt one of those rhythms, slow oscillation in particular, and actively contribute to Alzheimer’s progression. We tested the degree to which transplantation of healthy donor interneuron progenitors would restore slow oscillation rhythm in young APP mice. We harvested medial ganglionic eminence (MGE) progenitors from mouse embryos and transplanted them into host APP mutant cortices. 3D light-sheet and structured illumination microscopy revealed that transplanted MGE progenitors survived and matured into healthy interneurons. *In vivo* multiphoton calcium imaging and voltage-sensitive dye imaging showed functional integration and slow oscillation rescue in absence or presence of optogenetic stimulation. Our work provides proof-of-concept evidence that stem cell therapy may serve as a viable strategy to rescue functional impairments in cortical circuits of APP mice and potentially those of Alzheimer’s patients.

## Introduction

Alzheimer’s disease (AD) is a progressive neurodegenerative disorder that impairs cognitive functions, with aging as its greatest risk factor ^1^. Hallmark neuropathological features of AD include deposition of extracellular amyloid-beta (Aβ) plaques, presence of intracellular neurofibrillary tangles, synaptic dysfunction, and neurodegeneration in later stages ^2,3^. Recent therapeutic developments include U.S. Food and Drug Administration (FDA) approval of monoclonal antibodies targeting Aβ ^4,5^, such as Lecanemab and Donanemab, which reduce amyloid burden and modestly slow cognitive decline in some patients ^6,7^. However, individual responses vary ^4^. Concerns about adverse and potentially life-threatening events persist ^8^. Efficacy data in racially and ethnically diverse populations are limited. Finally, the high treatment costs of these therapies may restrict access ^5^. These constraints underscore the need for alternative or complementary therapeutic approaches aimed at additional pathological pathways to slow AD ^9^.

Alzheimer’s patients frequently report sleep impairments, which contribute to their disease progression ^10^. Aβ accumulations can further impair sleep, creating a positive feed-back relationship that exacerbates disease severity ^10–13^. Individuals at early stages of AD and mild cognitive impairment (MCI) consistently exhibit reduced non-rapid eye movement (NREM) sleep and impaired sleep-dependent brain rhythms, slow oscillation specifically ^14–16^. Because slow oscillation (low-frequency brain rhythm <1 Hz) is essential for synaptic plasticity and memory consolidation, its disruption accelerates memory decline during AD progression ^14,17^. Sleep disturbances can manifest at early stages of AD even preceding notable cognitive deficits. Consistent with clinical findings, APP/PS1 (APP) transgenic mice, a well-established model of amyloidosis, display reduced NREM sleep durations, and impaired slow oscillation ^18–21^. Hyperexcitability due to diminished inhibitory tone within cortical circuits underlies slow oscillation impairments in young APP mice ^18,19^. GABAergic interneurons balance neuronal hyperexcitability, maintain network homeostasis, and shape sleep architecture ^22,23^. Deficits in GABA signaling contribute to sleep impairment in AD ^24–26^. Furthermore, we found that optogenetic activation of endogenous cortical GABAergic interneurons restored NREM sleep, enhanced slow oscillation rhythm, slowed AD progression, and rescued sleep-dependent memory consolidation in APP mice ^19^. Thus, potentiating inhibitory tone during NREM sleep could ameliorate sleep impairments and potentially slow AD progression ^27^. Therefore, therapeutic strategies augmenting GABAergic interneuron function to restore slow oscillation early during disease progression are warranted.

Stem cell therapies are being pursued in the clinic for a variety of neurodegenerative diseases ^28–30^. Stem cell therapy holds promise as a treatment for AD ^31,32^. However, it remains unclear whether this approach can rescue cortical circuit function and slow oscillation deficits in AD. Would a single delivery of autologous cells that can engraft into local brain circuits and develop into the neurons of interest slow Alzheimer’s progression? Here, we tested whether a single delivery of autologous cells would engraft to the site of injury, develop into appropriate types of GABAergic neurons, and slow Alzheimer’s progression by restoring slow oscillation in APP mice. We harvested mouse medial ganglionic eminence (MGE) GABAergic cortical interneuron progenitors ^33–37^. The MGE is the birthplace of neocortical and hippocampal GABAergic interneurons. When transplanted into the host brain, MGE donor cells differentiated into functional inhibitory interneurons, restoring a healthy balance between excitatory and inhibitory neurotransmission ^33–35^. Thus, transplantation of MGE donor cells may alleviate AD-like phenotypes in mouse models ^36,37^.

In this study, fetal-derived MGE progenitors were harvested and then transplanted into adult host APP cortices. Donor cell survival and migration were assessed by 3D whole-brain lightsheet microscopy following tissue clearing. The fates and maturation of the transplanted cells were evaluated using established interneuron markers. Super-resolution structured illumination microscopy (SIM) was employed to investigate donor cell integration into host neural circuits. In vivo multiphoton microscopy was performed to monitor calcium transients using GCaMP6f targeted to donor cells, assessing their function within host circuits. Finally, voltage-sensitive dye (VSD) imaging was used to track slow oscillation in the absence and presence of optogenetic boost in APP mice. Collectively, these approaches confirm robust donor cell integration and provide insight into how MGE interneuron progenitor transplantation may alleviate network deficits associated with APP pathology. We demonstrated robust donor cell survival and synaptic integration in APP cortices, highlighting the therapeutic potential of MGE progenitor transplantation for Alzheimer’s disease.

## Results

### Transplantation of MGE Donor Progenitors into APP Hosts

Earlier studies reported that diminished inhibitory tone contributed to slow oscillation deficits in young APP mice ^18,19^. To assess the degree to which MGE interneuron progenitors restored inhibition and sleep-dependent brain rhythms, slow oscillation specifically, we transplanted them into B6C3 Tg(APPswe, PSEN1dE9)85Dbo/Mmjax ^38^ (APP) mice (Fig. 1). MGE progenitors were harvested from mouse embryos at embryonic day 13.5 (E13.5), the peak period for generating the cortical interneurons ^39^. Donor strains included VGAT-Venus, GP5.17, VGAT-ChR2-eYFP, and VGAT-Cre; Ai214. Two-month-old APP mice received a single injection of 500,000 MGE progenitor cells into the left anterior cortex (layers 2–5). Over the following two months, we evaluated the migration and maturation of Venus-expressing donor interneurons using histological analyses. We then monitored calcium transients in GCaMP6f-expressing donor cells *in vivo* to assess their function in the host brain circuit. Finally, we performed VSD imaging, in absence or presence, of light stimulation of ChR2- or GtACR1-expressing donor cells to determine whether donor neurons were necessary and sufficient to rescue slow oscillation.

**Fig. 1.**
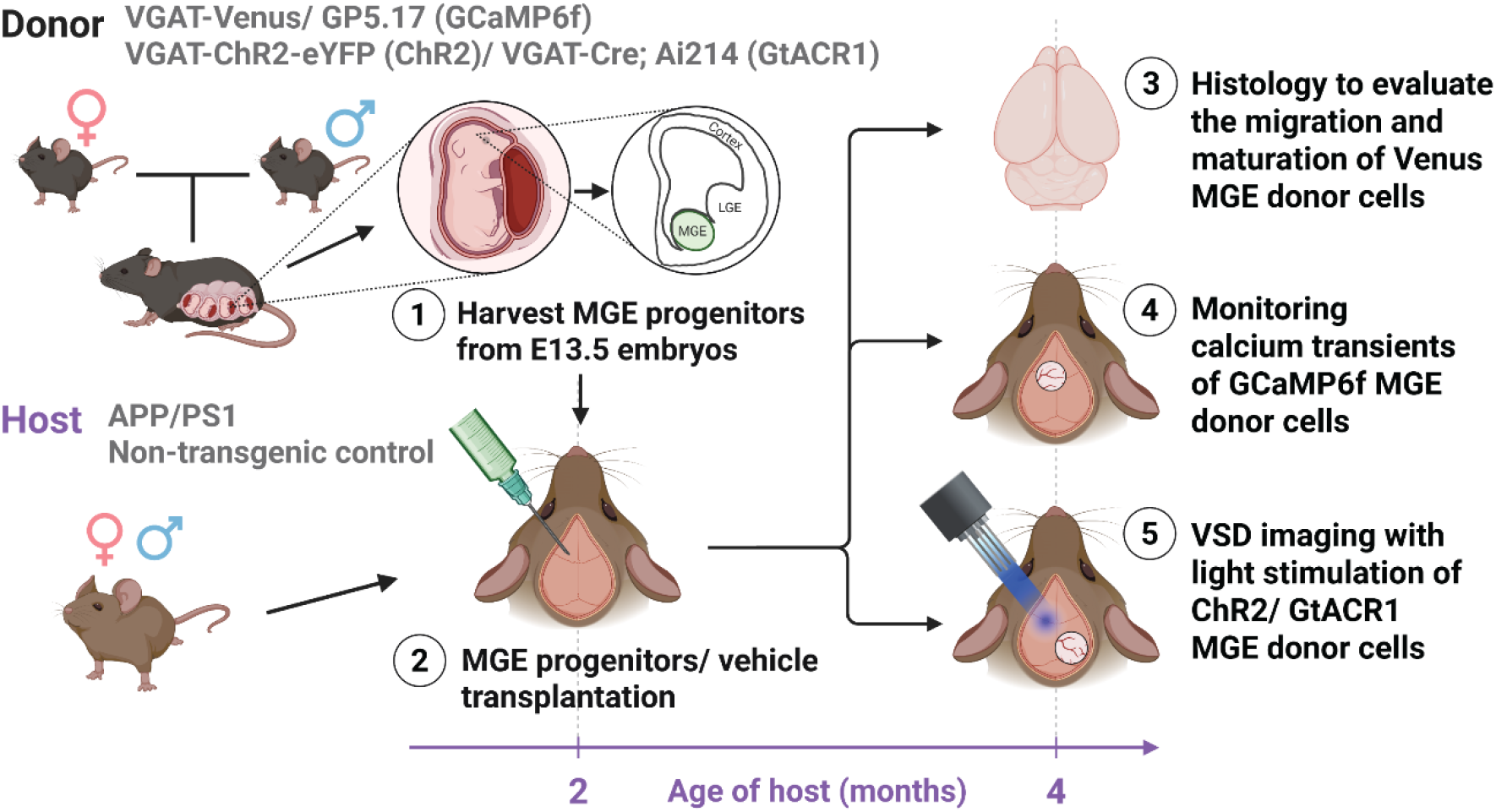
Study design. Donor strains (VGAT-Venus, GP5.17 [GCaMP6f], VGAT-ChR2-eYFP [ChR2], and VGAT-Cre; Ai214 [GtACR1]) were used to harvest medial ganglionic eminence (MGE) progenitors from mouse embryos on embryonic day 13.5 (E13.5). These progenitors were transplanted into the left anterior cortex (layers 2–5) of 2-month-old APP host mice. The donor-derived MGE cells were evaluated via histology, calcium transient monitoring (in GCaMP6f MGE donor cells), and voltage-sensitive dye (VSD) imaging with optogenetic stimulation (ChR2 or GtACR1 MGE donor cells) two months post-transplantation.

### Transplanted MGE Donor Progenitors Migrated within APP Host

We transplanted Venus-expressing MGE donor progenitors into APP brains to examine the survival of transplanted MGE donor cells and their migration in APP hosts. Two months post-transplantation, tissue clearing followed by whole-brain imaging using light-sheet microscopy revealed MGE donor cells in the anterior cortical regions (Fig. 2a, Supplemental Video 1), confirming their survival. Higher magnification views show that MGE donor cells migrated beyond the injection site within the host cortices (Fig. 2b, c). On average, 1,118 ± 334 migrant MGE donor cells were detected per cortex (Fig. 2d), corresponding to 0.26 ± 0.024% compared to the total MGE cells injected (Fig. 2e). The mean migratory distance was 0.63 ± 0.39 mm (Fig. 2f), with 94.3% of cells migrating within 1 mm of the injection site. Few donor cells were detected as far as 5 mm (Fig. 2g). Over the two-month survival period that was examined in this study, the migration of donor cells appeared to be relatively limited within the site of transplantation in the anterior cortex. In addition, we observed no significant differences in cell survival between non-transgenic and APP mice (Supplemental Fig. 1), suggesting that the cortical environment in APP mice does not negatively affect MGE donor cell viability. Overall, our results indicate that transplanted MGE cells survived and showed limited migration within the host cortices.

**Fig. 2.**
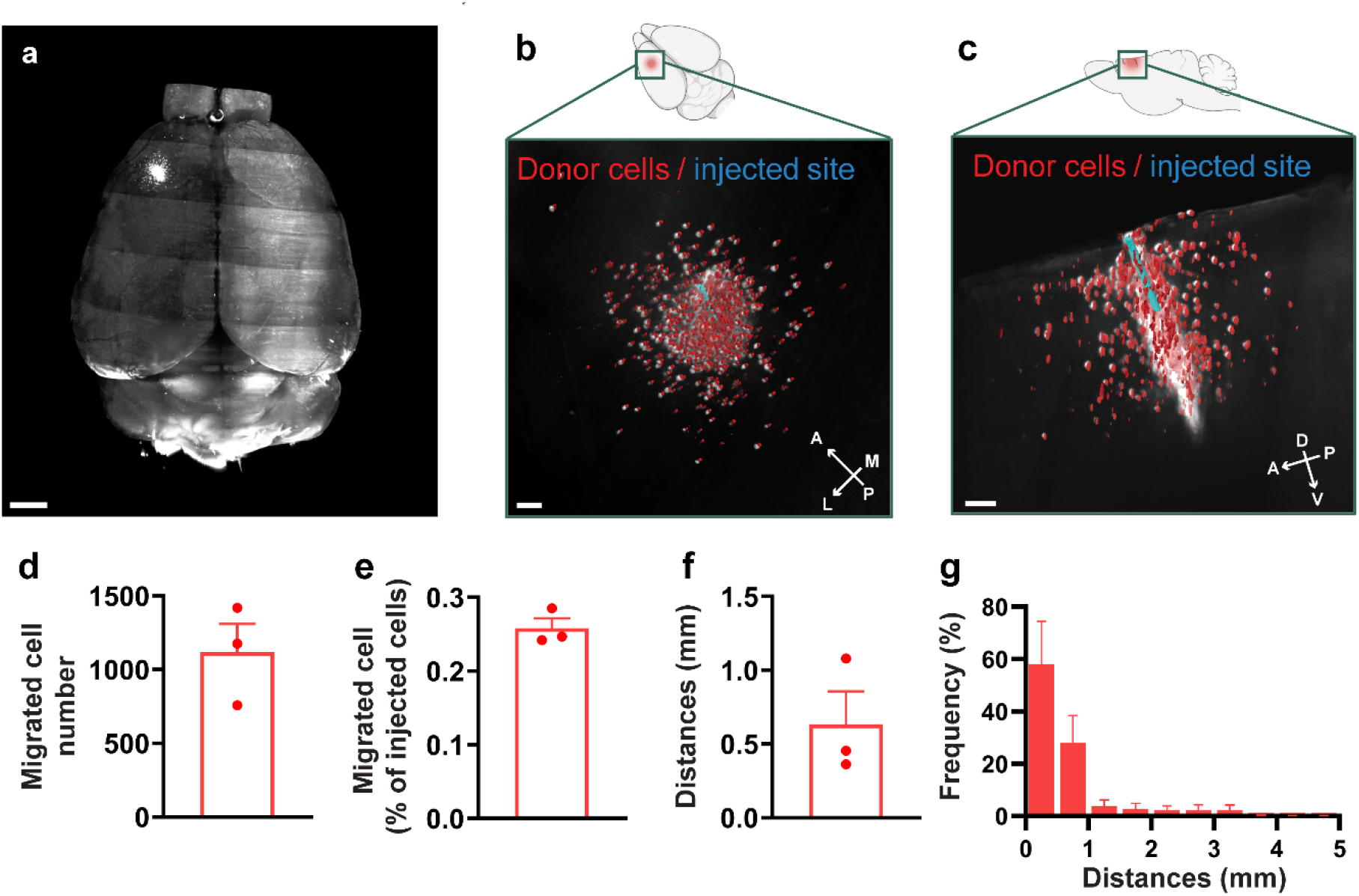
MGE donor cells transplanted into the APP host cortices survived and migrated for 60 days. **a** 3D reconstruction of the host whole brain. **b, c** Higher-magnification 3D reconstructions from the dorsal (**b**) and sagittal (**c**) views, showing donor cells (red) and the injection site (blue). **d** Number of migrated cells detected by whole-brain imaging. **e** Percentage of migrated cells, calculated as the number of migrated cells divided by the total number of transplanted cells. **f** Average migration distance, defined as the distance between donor cells and the injection site. **g** Distribution of migration distances. Data are shown as mean ± SEM. Scale bars, 1 mm (**a**), 0.2 mm (**b**), and 0.3 mm (**c**). *n* = 3 mice, biologically independent replicates.

### MGE Donor Progenitors Matured into Healthy Interneurons

To investigate whether transplanted MGE progenitors differentiated into mature interneurons in APP mice, we transplanted Venus-expressing MGE donor cells into the host cortices (Fig. 3a). We evaluated cell fate two months post-transplantation using immunohistochemistry. First, we verified that donor cells localized to the same cortical regions observed with whole-brain imaging (Fig. 2a, 3b). The transplanted donor cells expressed little to no astrocyte (GFAP), microglial (Iba1) or oligodendrocyte (Olig2) markers (Fig. 3c-f). The MGE donor cells showed robust expression of the neuronal marker NeuN (Fig. 3c–f; 0.20 ± 0.55% GFAP [GFAP vs. NeuN; *p* = 0.0007], 0.44 ± 0.89% Iba1, [Iba1 vs. NeuN; *p* = 0.0019] 0.29 ± 0.88% Olig2 [Olig2 vs. NeuN; *p* = 0.0006], 77.0 ± 5.7% NeuN). Consistent with a GABAergic interneuron phenotype, most transplanted cells were GAD67-positive (85.1 ± 7.5%) but lacked CaMKII expression (0.56 ± 1.2%, [GAD67 vs. CaMKII; *p* = 0.0079]), confirming their GABAergic interneuron identity (Fig. 3g–i).

**Fig. 3.**
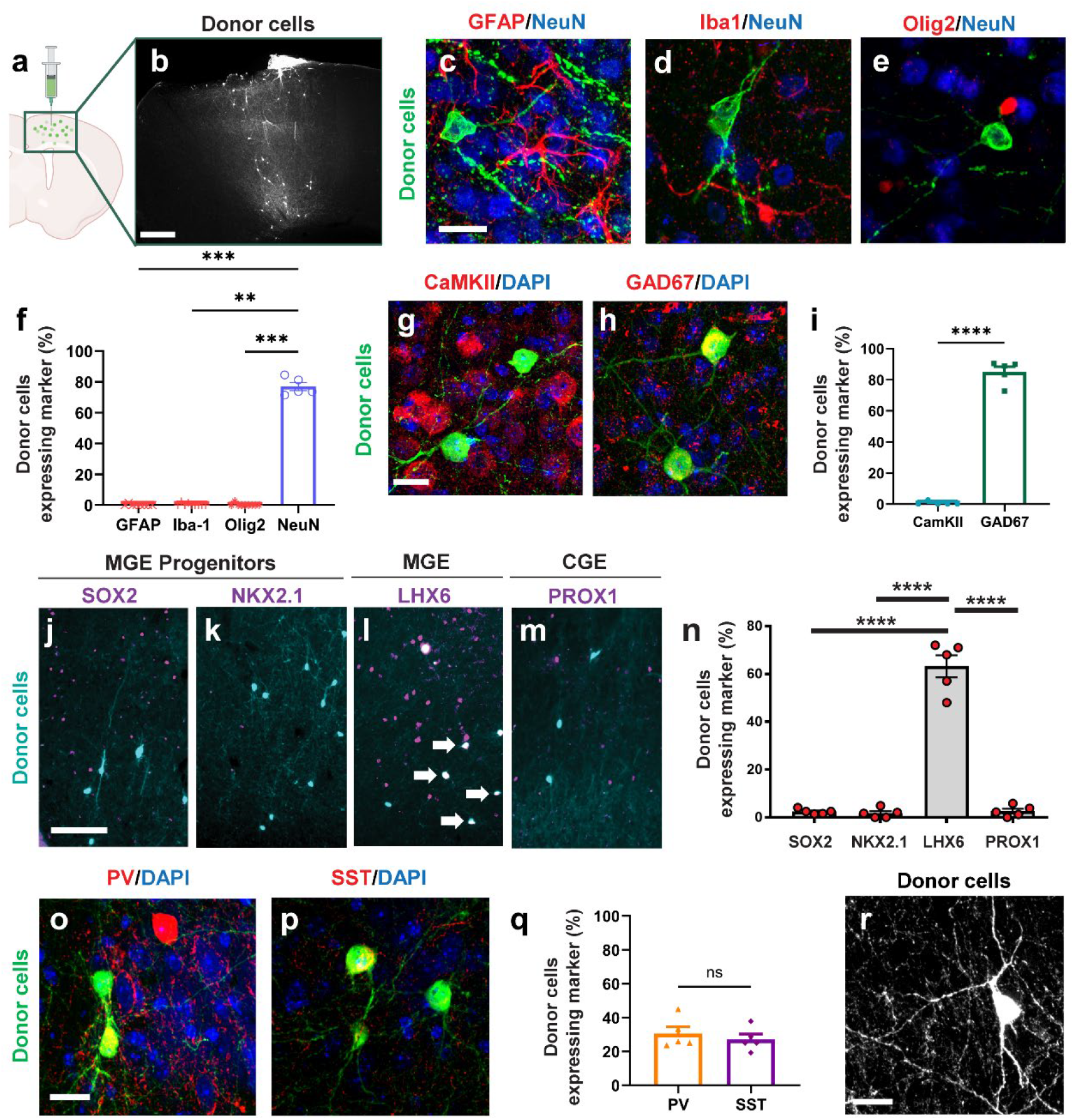
MGE donors transplanted into the APP hosts matured into interneurons over 60 days. **a** Schematic of medial ganglionic eminence (MGE) donor cell transplantation. **b** MGE donor cell distribution 2 months post-transplantation. Donor cells were transplanted into the host anterior neocortex and visualized following immunostaining with anti-GFP antibody. **c–e** Immunolabeling of the proportion neuronal and glial markers in the host anterior neocortex (GFAP, Iba-1, Olig2, NeuN). **f** Quantification of donor cell co-labeling with neuronal and glial markers. Statistical analysis was conducted using Kruskal– Wallis test followed by Dunn’s multiple comparisons test (GFAP vs. NeuN; *p* = ^***^0.0007, Iba1 vs. NeuN; ^*\*\**^*p* = 0.0019, Olig2 vs. NeuN; ^***^*p* = 0.0006). **g, h** Immunolabeling of excitatory (CaMKII) and inhibitory (GAD67) neuron markers in the host anterior neocortex, with DAPI. **i** Quantification of the proportion of donor cell co-labeling with excitatory and inhibitory markers. Statistical analysis is conducted using Kruskal–Wallis test followed by Mann-Whitney *U* Test test (GAD67 vs. CaMKII; *\*\*p* = 0.0079). **j–m** Immunolabeling of progenitor/proliferating cells (SOX2, NKX2.1), MGE-derived cells (LHX6), and CGE-derived cells (PROX1) markers in the host anterior neocortex. **n** Quantification of the proportion of donor cells expressing maturation markers. Statistical analysis was conducted using Kruskal–Wallis test followed by Dunn’s multiple comparisons test (SOX2 vs. LHX6; *p* = xx, NKX2.1 vs. LHX6; *p* = xx, PROX1 vs. LHX6; *p* = xx). **o, p** Immunolabeling of mature MGE interneuron markers (SST, PV) with DAPI in host anterior neocortex. **q** Quantification of the proportion of donor cell co-labeling with mature MGE interneuron markers. Statistical analysis was conducted using Student’s *t*-test (PV vs. SST; *p* = 0.51). **r** Representative Z-projection confocal images (average intensity) of donor cells. Data are presented as mean ± SEM (**f, i, q, n**). ns, not significant; ^*^*p* < 0.05, ^**^*p* < 0.01, ^***^*p* < 0.001, ^****^*p* < 0.0001. Scale bars: 400 µm (**a**), 20 µm (**c, g, o, r**), and 100 µm (**j**). *n* = 5 mice, biologically independent replicates.

We next examined maturation. We found that the MGE donor cells expressed little to no progenitor markers SOX2 (2.42 ± 0.48%, [SOX2 vs. LHX6; *p* < 0.0001]) or NKX2.1 (1.74 ± 0.9%, [NKX2.1 vs. LHX6; *p* < 0.0001]). The donor cells expressed LHX6 (63.2 ± 4.6%, Fig. 3j–l, n), consistent with an MGE interneuron lineage ^40,41^. The donor cells expressed little to no CGE-derived marker PROX1 (Fig. 3m, n, 2.6 ± 1.01%, [PROX1 vs. LHX6; *p* < 0.0001]). Staining for NKX2.1 in the striatum (Supplemental Fig. 2o) served as a positive control since NKX2.1 expression ceases once MGE-lineage interneurons enter the neocortex but persists in striatal interneurons, which derive from a different lineage. Subsets of MGE donor cells expressed parvalbumin (PV; 30.9 ± 8.9%, [PV vs. SST; *p* = 0.51]) or somatostatin (SST; 27.2 ± 6.9%), verifying their interneuron subtypes (Fig. 3o–q) ^34^. Morphologically, MGE donor cells displayed characteristic interneuron features, such as complex dendritic architecture and large soma size (Fig. 3r). Additional immunostaining revealed the presence of endogenous GFAP-positive astrocytes, but not microglia, near the injection site (Supplemental Fig. 2a–l). Importantly, the lack of immunostaining for a marker of proliferation, Ki67 confirmed that after transplantation, MGE donor cells were post-mitotic and non-tumorigenic (0.0%, *n = 5* mice. Supplemental Fig. 3m). The presence of Ki67 signal was verified within lymph node tissue, which is rich in proliferating cells (Supplemental Fig. 2n). Overall, these findings demonstrated that transplanted MGE progenitors successfully differentiated into mature GABAergic interneurons with lineage-appropriate identity and did not form tumors in the APP host cortices.

### Donor Interneurons Formed Synapses with APP Host Neurons

We used super-resolution structured illumination microscopy (SIM) to investigate whether the transplanted MGE interneurons integrated structurally into the host neural circuitry. We observed that Venus-expressing donor interneurons received both putative excitatory and inhibitory inputs and made putative inhibitory synapses targeting other neurons. Putative inhibitory synapses were identified using the presynaptic markers Bassoon and VGAT, and the postsynaptic marker gephyrin (Fig. 4a, f). Putative excitatory synapses were marked by the presynaptic marker Bassoon and the postsynaptic marker PSD95 (Fig. 4k). We observed that axonal boutons of the donor interneurons made inhibitory synapses targeting unlabeled host neurons (Fig. 4b–e). In addition, dendrite-like processes of the donor interneurons received both inhibitory (Fig. 4g–j) and excitatory inputs (Fig. 4l–o). Quantitative analyses showed that donor axon-like processes exhibited 2.4 ± 1.0 inhibitory synapses per 10 µm of donor process (inhibitory presynapses vs. inhibitory postsynapses; *p* = 0.016, inhibitory presynapses vs. excitatory postsynapses; *p* = 0.88) while dendrite-like processes contained 0.99 ± 0.6 inhibitory synapses and 2.2 ± 0.42 excitatory synapses per 10 µm (Fig. 4p, inhibitory postsynapses vs. excitatory postsynapses; *p* = 0.043). These synaptic densities approximated those in healthy nontransgenic neurons ^42–44^ Thus, our results indicate that the transplanted donor interneurons exhibited structural inhibitory and excitatory synapses with host neurons, suggesting that the transplanted interneurons establish synaptic connections within the host brains and integrate into the host neural networks.

**Fig. 4.**
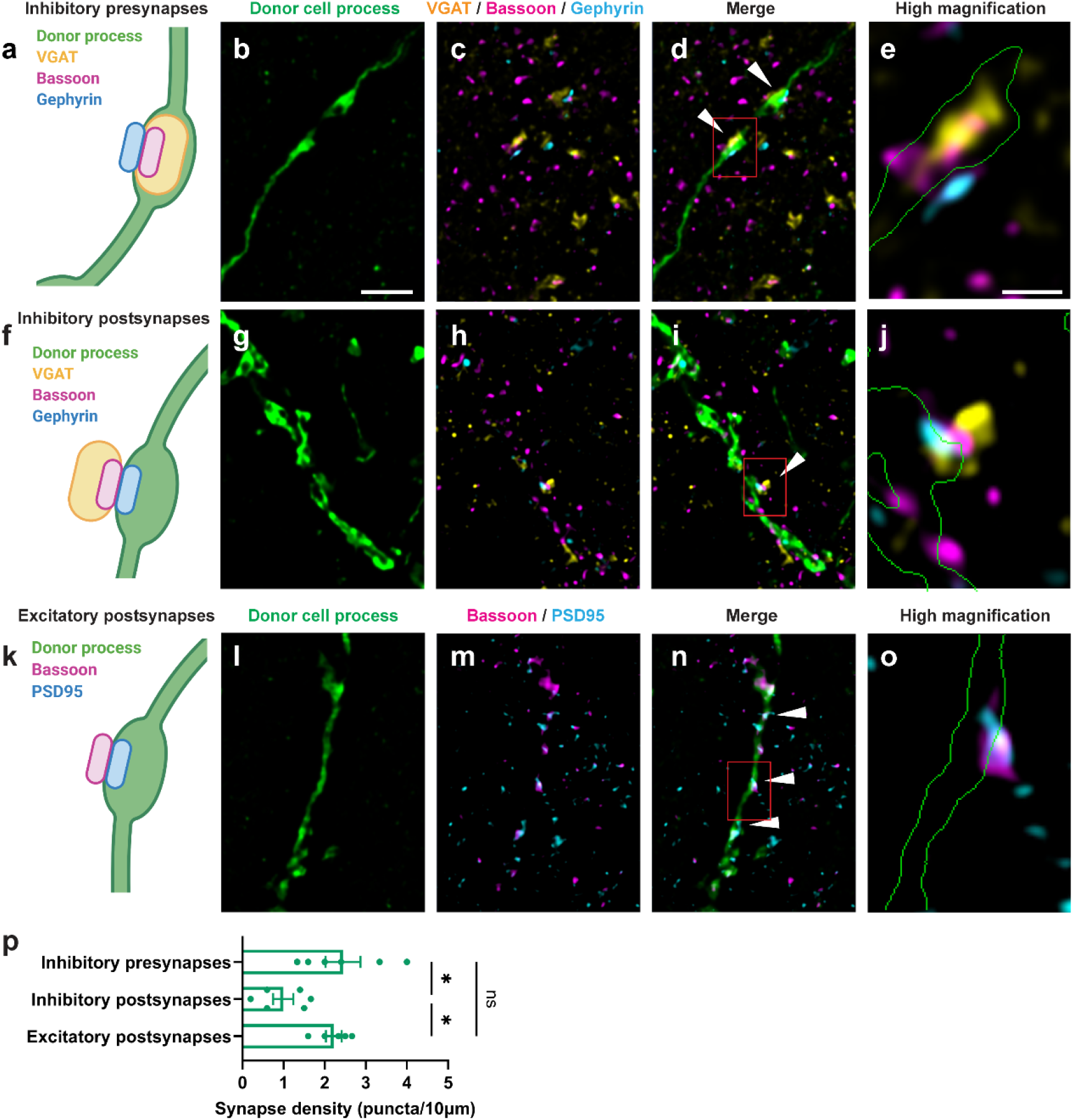
MGE donor cells form synaptic connections with APP host neurons. **a** Schematic illustrating inhibitory presynapses in donor axon-like processes. b–d Super-resolution structured illumination microscopy (SIM) images of synaptic marker labeling (VGAT, Bassoon, Gephyrin) in donor axon-like processes in the host anterior neocortex. **e** Higher-magnification SIM image of inhibitory presynapse in donor axon-like processes in the host anterior neocortex with the outline of donor cell process (green). **f** Schematic illustrating inhibitory postsynapses in donor dendrite-like processes. **g–i** SIM images of synaptic marker labeling (VGAT, Bassoon, Gephyrin) in donor dendrite-like processes in the host anterior neocortex. **j** Higher-magnification SIM image of inhibitory postsynapse in donor dendrite-like processes in the host anterior neocortex with the outline of donor cell process (green). **k** Schematic illustrating excitatory postsynapses in donor dendrite-like processes. **l–n** SIM images of synaptic marker labeling (VGAT, Bassoon, PSD95) in donor dendrite-like processes. **o** Higher-magnification SIM image of excitatory postsynapse in donor dendrite-like processes in the host anterior neocortex with the outline of donor cell process (green). **p** Quantification of synaptic density (per length of donor process). The white arrowheads indicate synapses. The red rectangles in figures d, i, and n indicate the enlarged areas in figures e, j, and o, respectively. Statistical analysis was conducted using the one-way ANOVA followed by Tukey’s multiple comparisons test (Inhibitory presynapses vs. Inhibitory postsynapses; ^*^*p* = 0.016, Inhibitory presynapses vs. Excitatory postsynapses; *p* = 0.88, Inhibitory postsynapses vs. Excitatory postsynapses; ^*^*p* = 0.043). Data are presented as mean ± SEM Ns, not significant; ^*^*p* < 0.05. Scale bar: 2 µm. *n* = 5–6 mice, biologically independent replicates.

### Donor Interneurons Were Incorporated into APP Host Circuits

To further investigate whether donor interneurons integrated functionally into the host neural circuitry, we transplanted GCaMP6f-expressing MGE progenitors from GP5.17 donor mice and monitored donor calcium dynamics via multiphoton microscopy through cranial windows, two months post-transplantation. Donor interneurons exhibited robust GCaMP6f signals (Fig. 5a). We detected calcium transients in donor interneurons (Fig. 5b). Quantification of event rates from ΔF/F traces yielded an average of 0.10 ± 0.073 Hz (Fig. 5c), consistent with previously reported cortical interneuron activity levels ^24^. Thus, we provide structural and functional evidence that donor interneurons integrate physiologically into the APP cortices.

**Fig. 5.**
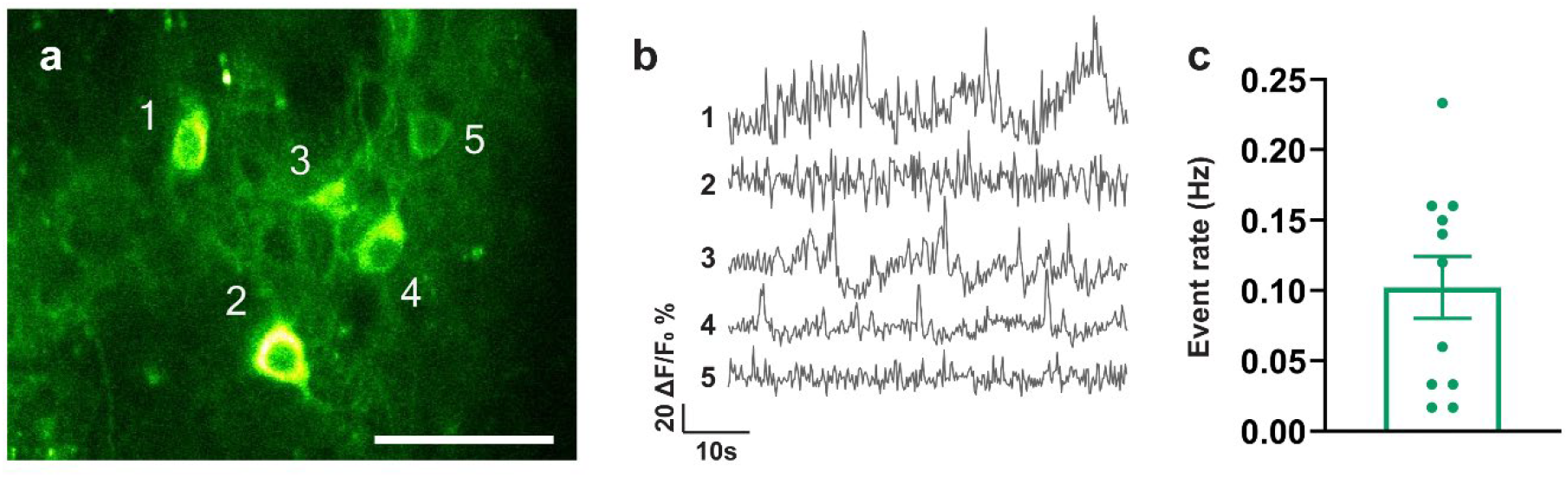
MGE donor cells exhibited calcium transients in the APP host cortices. **a** In vivo fluorescence images of GCaMP6f-labeled donor interneurons in the anterior cortex of APP mice. **b** Representative raw traces of donor calcium transients in the APP hosts. **c** Quantification of the spontaneous calcium event rates. Data are presented as mean ± SEM Scale bar: 50 µm. n = 11 neurons from 3 mice, biologically independent replicates.

### MGE Transplantation Restored Slow Oscillation

APP mice exhibited aberrant slow oscillation. Specifically, slow oscillation power was low. We therefore investigated the degree to which donor transplants could restore slow oscillation in the APP mouse cortex. We transplanted donor progenitors expressing VGAT-Venus, VGAT-ChR2-eYFP, VGAT-Cre; Ai214. Mice randomly assigned to Vehicle group were injected with cell-free intracranial transplantation media and served as negative controls. Cell transplants and Vehicle injections were made into the anterior left cortical hemispheres. Two months post-transplantation, voltage-sensitive dye (VSD) RH2080 imaging of contralateral right hemispheres (Fig. 6a) revealed a 0.5–1.0 Hz (peak ∼0.6 Hz) band of cortical slow oscillation. This frequency band was previously reported to be significantly lower in APP mice compared to non-transgenic wildtype controls ^18,20^. Regions of interest (ROIs) were defined in the VSD images, and the resulting ΔF/F traces (Fig. 6b, d, f, h) were subjected to Fourier analysis to quantify slow oscillation power (Fig. 6c, e, g, i). Donor-transplanted APP mice exhibited significant increases of slow oscillation power compared to vehicle-treated APP hosts (Fig. 6c, e, j, 0.22 ± 0.10 ×10^−7^ control in Vehicle vs. 1.1 ± 0.47 ×10^−7^ control in Venus-donor; *p* = 0.0033). Furthermore, optogenetic activation of ChR2 at the endogenous frequency of slow oscillation, 0.6 Hz, further increased slow oscillation power (Fig. 6g, j, 0.92 ± 0.73 ×10^−7^ control in ChR2-donor vs. 1.8 ± 0.73 ×10^−7^ light activation in ChR2-donor; *p* < 0.0001). The optogenetically-induced boost in slow oscillation power was absent in Venus-donor or vehicle-injected controls. These findings indicate that donor interneuron activation is sufficient to potentiate slow oscillatory activity. In contrast, inhibitory optogenetic stimulation of GtACR1 reduced slow oscillation power compared to no-light stimulation controls (Fig. 6i, k, 1.0 ± 0.39 ×10^−7^ control vs. 0.70 ± 0.37 ×10^−7^ light stimulation; *p* = 0.0094), partially reversing the transplantation-induced rescue. Thus, donor cell activity is required for restoring slow oscillation. In addition, GtACR1-inhibitory effect appeared specific to continuous-wave light stimulation, as random wave stimulation failed to suppress the slow oscillation rescue (Supplemental Fig. 3). Altogether, these results demonstrated that donor interneurons were necessary and sufficient to restore slow oscillation in the APP cortices. In conclusion, our findings highlight the therapeutic potential of MGE progenitor transplantation to restore slow oscillation in this mouse model of Alzheimer’s disease.

**Fig. 6.**
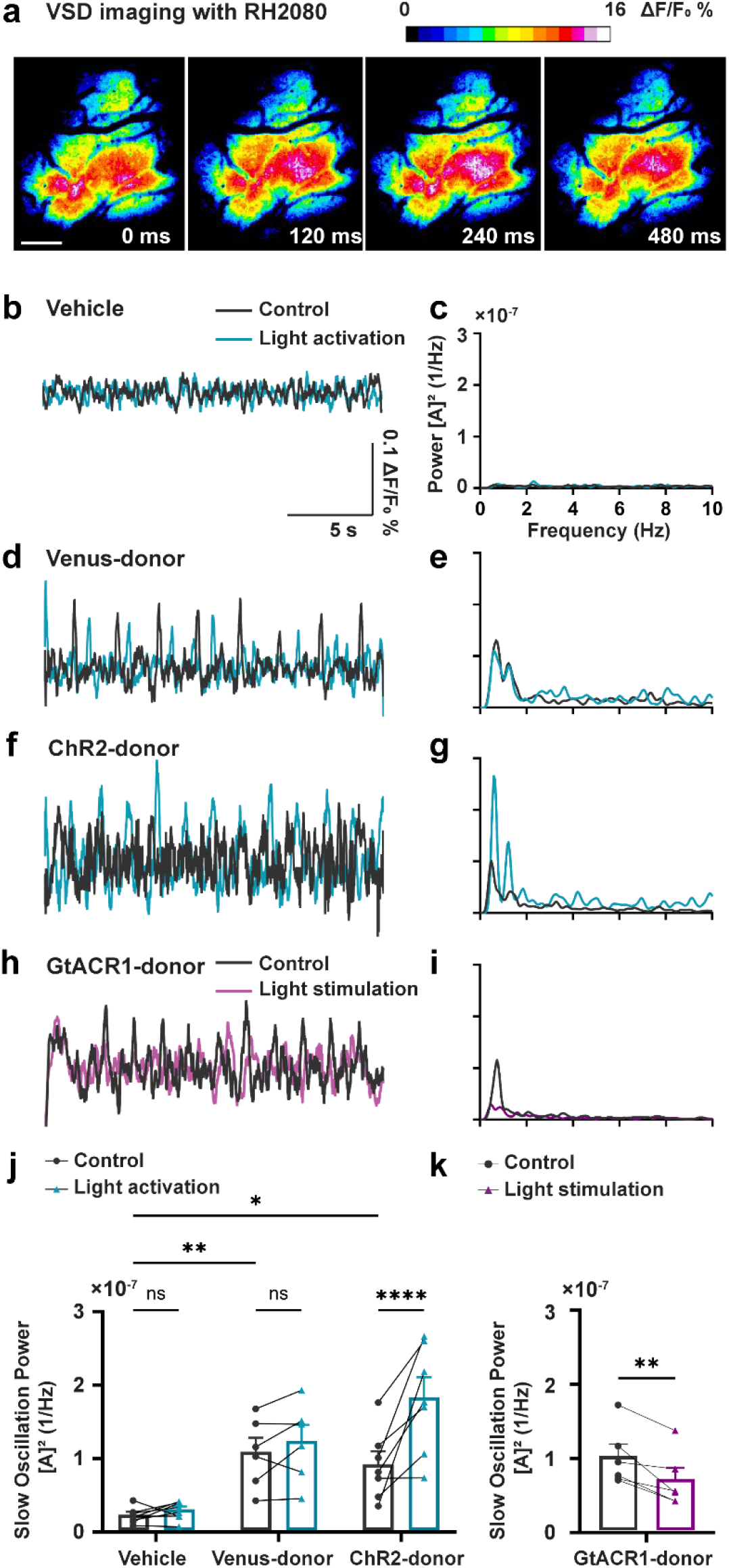
MGE cell transplantation rescued slow oscillation in APP mice. **a** Representative in vivo voltage-sensitive dye (VSD) RH2080 images of the somatosensory cortex in an APP mouse, showing oscillatory activity. Scale bar: 100 µm. **b** Representative raw fluorescence traces of VSD imaging in the vehicle group (transplantation medium without MGE donor cells) with (blue) or without (gray) 0.6 Hz pulse wave optogenetic stimulation 2 months after transplantation. **c** Representative power spectral density analysis of traces in the vehicle group with (blue) or without (gray) 0.6 Hz pulse wave optogenetic stimulation. [A] ^2^ = magnitude of the Fourier amplitude squared. **d** Representative raw fluorescence traces in the Venus-MGE group (MGE donor cells expressing VGAT-Venus as no optogenetic opsin control) with (blue) or without (gray) 0.6 Hz optogenetic stimulation 2 months after transplantation. **e** Representative power spectral density analysis of traces in the Venus-MGE group with (blue) or without (gray) 0.6 Hz pulse wave optogenetic stimulation. **f** Representative raw fluorescence traces in the ChR2-MGE group (MGE donor cells expressing VGAT-ChR2-eYFP as optogenetic activation of MGE donor cells) with (blue) or without (gray) 0.6 Hz optogenetic stimulation 2 months after transplantation. **g** Representative power spectral density analysis of traces in the ChR2-MGE group with (blue) or without (gray) 0.6 Hz pulse wave optogenetic stimulation. **h** Representative raw fluorescence traces in the GtACR1-MGE group (MGE donor cells expressing VGAT-Cre; Ai214 as optogenetic inhibition of MGE donor cells) with (purple) or without (gray) continuous wave optogenetic stimulation 2 months after transplantation. **i** Representative power spectral density analysis of traces in the GtACR1-MGE group with (purple) or without (gray) continuous wave optogenetic stimulation. **j** Slow oscillation (0.5–1.0 Hz) power with or without 0.6 Hz optogenetic activation. Each data point represents the average of 10–15 traces from each mouse. Statistical analysis was conducted using the repeated measure two-way ANOVA followed by Šídák’s multiple comparisons test (control in Vehicle vs. control in Venus-MGE; ^**^*p* = 0.0033, control in Vehicle vs. control in ChR2-MGE; ^*^*p* = 0.0165, control in Vehicle vs. light activation in Vehicle; *p* = 0.6371, ^*^*p* = 0.0165, control in Venus-MGE vs. light activation in Venus-MGE; *p* = 0.3814, control in ChR2-MGE vs. light activation in ChR2-MGE; ^****^*p* < 0.0001). **k** Slow oscillation (0.5–1.0 Hz) power with optogenetic stimulation. Each data point represents the average of 10–15 traces from each mouse. Statistical analysis is conducted using the Paired Student’s *t*-test (control vs. light stimulation; ^**^*p* = 0.0094). Data are presented as mean ± SEM Ns, not significant; ^*^*p* < 0.05, ^**^*p* < 0.01, ^***^*p* < 0.001, ^****^*p* < 0.0001. *n* = 5–7 mice, biologically independent replicates.

## Discussion

Our overall goal was to determine the degree to which transplantation of MGE interneuron progenitors into an APP mouse model of AD could restore slow oscillation. Soluble oligomeric Aβ is detected in 4-month-old APP mice, but amyloid plaque deposition is not yet observed. Our study demonstrates that transplanting MGE interneuron progenitors that mature into mature interneurons in 4-month old APP mice restores slow oscillation. Importantly, MGE donor cells successfully survived and migrated within the host cortices, differentiated into mature GABAergic interneurons, and established functional inhibitory circuits. These results indicate that enhancing inhibitory tone via MGE-derived interneurons can mitigate sleep-dependent brain rhythm impairments and neuronal network dysfunction.

Consistent with previous findings using serial coronal sections ^33,45,46^, our 3D tissue clearing and light-sheet microscopy revealed robust survival and migration of MGE donor cells in APP cortex, providing a more comprehensive spatial perspective compared to 2D conventional histological approaches. Although many transplanted cells stayed within the anterior left cortex in close proximity to the injection site, VSD imaging showed improved slow oscillation power throughout the contralateral somatosensory cortex, suggesting that restoring local inhibition can influence the broader host cortical network. We targeted the prefrontal cortex for transplantation because the prefrontal cortex is the site of intrinsic generation of slow oscillation during sleep ^47–50^. Interneurons are known for synchronizing slow oscillatory activity ^51,52^, and it is thought that a small number (on the order of only a few hundred cells) of cortical neurons can initiate or maintain slow oscillatory activity ^22,53,54^. Our results indicated that the donor interneurons in the prefrontal cortex could facilitate the propagation of slow oscillation to the contralateral somatosensory cortex. However, whether MGE-derived neurons extended long-ranging axonal projections or primarily modulated local circuits remain unknown. Notably, Henderson and colleagues ^55^ reported that MGE donor cells transplanted into the dentate gyrus can extend axons across the hippocampal commissure or into the medial entorhinal cortex in other mouse models, suggesting that the transplanted cells could influence oscillatory dynamics through extensive axonal projections to distant cortical regions. Future experiments using transsynaptic tracing are necessary to determine whether that is possible in AD mouse models.

Despite successful transplantation, the overall survival rate of MGE donor cells was relatively low compared with previous MGE transplantation studies ^56,57^, potentially due to practical constraints such as the competition of the large number of transplanted cells for limited neurotrophic support and limited host cortical capacity. This experiment involves transplanting MGE donor cells into the cortex. Compared to earlier studies involving transplantation of cells into the hippocampus or other deep regions, we injected cells into the neocortex, which is prone to leakage during transplantation, that can affect survival rates. We transplanted 500,000 cells per procedure which is greater than the amounts typically reported in similar experiments ^56,57^. Notably, no significant differences in cell survival were detected between non-transgenic and APP mice (Supplemental Fig. 1), suggesting that the cortical environment of APP mice is unlikely to negatively impact MGE donor cell viability. In summary, these data demonstrate that MGE donor cells can survive and migrate effectively in the APP mutant cortex.

We verified that MGE donor cells differentiated into mature GABAergic interneurons with established MGE lineage subtypes ^33,54^. SOX2 and NKX2.1 are important for establishment of the developmental trajectory of MGE-derived lineages but are nearly absent in mature neocortical interneurons ^46^. LHX6 is essential for the generation of SST and PV cortical interneurons within the MGE lineage. It is also required for their migration to the cortex and functions as a transcription factor in directing cell fate^40,58,59^. PROX1 serves as a CGE lineage marker. Absence of PROX1-positive cells confirmed that the transplanted cells originated from the MGE. Observing significant maturation at two months post-transplantation aligns with previous research on MGE progenitors ^34^. Furthermore, the restricted expression of PV and SST in a subset of MGE donor cells is consistent with prior findings ^55,60^. Taken together, these data confirm that transplanted progenitors mature into healthy MGE-derived interneurons in the host cortex. In addition, our Ki67 staining indicates no tumorigenesis in MGE donor cells, consistent with previous reports ^45^. Tumorigenesis is a major concern during stem cell therapy ^61^. On the other hand, we observed GFAP-positive areas surrounding the injection sites, suggesting that tissue damage or cell debris during transplantation may induce chronic astrogliosis.

We demonstrated that MGE donor cells establish inhibitory synapses along axon-like processes in host tissue. This observation aligns with other results that MGE donor cells expressing LHX6 can differentiate into SST- or PV-positive interneurons, indicating the developmental readiness to form synaptic connections. Our results also agree with published work showing synaptophysin, VGAT, and gephyrin co-localization within MGE donor cells ^55,62,63^. Gupta and colleagues also demonstrated that donor interneurons formed functional inhibitory synapses with host neurons using patch-clamp electrophysiology ^64^. As part of Alzheimer’s progression, impaired inhibitory interneuron function reduces inhibitory tone, contributing to hyperexcitation and network dysfunction ^13,25^. Our earlier studies revealed lower cortical expression of GABA, as well as GABA_A_ and GABA_B_ receptors, in APP mice ^18^. Topical GABA administration and optogenetic stimulation of endogenous GABAergic interneurons restored slow oscillation in brains of APP mice ^19^. Consequently, we suggest that MGE transplantation restores slow oscillation via healthy synaptic connectivity and GABAergic signaling, which could serve as a one-time and potentially permanent therapy. Previous AD mouse model research supports this mechanism, showing that MGE-derived interneurons re-establish circuit functions by forming inhibitory synapses. For example, Tong and colleagues ^36^ found that embryonic MGE-derived progenitors transplanted into ApoE4 knock-in mice boosted GABAergic inhibitory currents, restored excitatory/inhibitory balance, and improved learning and memory. Similarly, Lu and colleagues ^37^ reported that transplanting embryonic MGE progenitors into dentate gyrus of 7-month-old APP mice led to differentiation into GABAergic subtypes. This process suppressed hippocampal hyperexcitability, enhanced synaptic plasticity, and ultimately rescued cognitive deficits.

In addition to anatomical incorporation, our findings underscore the functional integration of MGE donor cells into host neural networks ^33^. Using GCaMP6f, we detected calcium transients in MGE donor cells, indicating active participation in host circuits. These results are consistent with earlier studies demonstrating intrinsic firing properties in transplanted MGE cells ^33,65^. Although we analyzed a limited number of cells, these cells could be subdivided into two distinct groups based on their firing rates. These likely correspond to SST-positive and PV-positive interneurons since SST cells usually fire at lower frequencies compared to PV cells ^24^. Future studies are needed to characterize the firing properties of donor interneurons using electrophysiological methods and determine the full extent of their identities. Consistent with immunostaining results, which confirm the presence of SST-positive and PV-positive neurons, these calcium imaging data support the conclusion that MGE donor cells mature into healthy interneurons and integrate into host circuits.

Our VSD experiments revealed that transplanted MGE donor cells significantly increase slow oscillation power. Light stimulation of ChR2 further increases slow wave power, while light activation of GtACR1 decreases slow wave power. These observations highlight that MGE donor cells are necessary and sufficient to restore slow oscillation. We previously reported that optogenetic stimulation of endogenous GABAergic interneurons restored slow oscillation power during NREM sleep in APP mice ^19^. Here, an analogous approach targeting exogenous neurons yielded a similar rescue. These findings extend prior research linking weakened GABAergic function to sleep deficits in AD. They also reinforce the broader concept that targeting neuronal circuits can address core neurophysiological processes in AD. Since slow oscillation rhythm during NREM sleep is important for memory consolidation and glymphatic clearance ^13^, potentiating inhibition and thus restoring inhibition/excitation balance via MGE interneurons could alleviate sleep impairments and reduce cognitive decline, as well as amyloid pathology.

Despite these promising results, several questions remain. Here, we focused on slow oscillation in 4-month-old APP mice, but it is unclear whether MGE transplantation can rescue sleep and slow Alzheimer’s progression in older APP mice. In previous work, optogenetic activation of endogenous interneurons improved slow oscillation, rescued sleep, slowed amyloid deposition, and restored memory functions ^19^, suggesting that MGE transplantation may yield comparable benefits. While no tumorigenesis or serious adverse reactions were observed during the two-month window, the long-term safety and stability of MGE donor cells requires further study. Future work should also investigate whether human stem cell-derived interneurons ^66–70^ can achieve similar outcomes and how best to refine transplantation protocols for clinical settings. It will be important to determine whether enhanced slow oscillation translate into sustained cognitive and neuropathological improvements over extended periods and across different AD models. Moreover, because impaired inhibitory function in AD may overlap with other disease mechanisms, research is needed to clarify how MGE transplantation interacts with these processes. Overcoming these challenges could position stem cell transplantation as a complementary option alongside existing treatments, such as monoclonal antibodies. Ultimately, cell-based therapies may enable fundamental circuit-level repair that improves sleep quality, cognitive performance, and clinical outcomes, affirming MGE transplantation’s promise as an alternative or adjunct to amyloid-focused approaches.

In summary, we show that MGE progenitor transplantation can restore sleep-related circuit function in an AD mouse model. The transplanted MGE donor cells differentiate into mature interneurons, reestablish inhibitory tone, and restore slow oscillation, which plays an important role in memory consolidation. Our findings suggest that stem cell therapy aimed at restoring neural circuits may offer a promising approach to improving sleep-dependent brain rhythms and slowing AD progression.

## Methods

### Animals

C57BL/6J mice (Jackson stock# 000664), VGAT-ChR2-EYFP mice (B6.Cg-Tg(Slc32a1 COP4^*^H134R/EYFP)8Gfng/J; Jackson stock #014548), VGAT-Cre mice (B6J.129S6(FVB)-Slc32a1tm2(cre)Lowl/MwarJ; Jackson stock #028862), Ai214 mice (B6.Cg-Igs7tm214(CAG-ACR1^*^, CAG-mRuby3)Tasic/J); Jackson stock #037380), and GP5.17 mice (C57BL/6J-Tg(Thy1-GCaMP6f)GP5.17Dkim/J; Jackson stock #025393) were purchased from Jackson Laboratories (Bar Harbor, USA). B6C3 Tg(APPswe, PSEN1dE9)85Dbo/Mmjax, RRID: MMRRC_034829-JAX, was obtained from the Mutant Mouse Resource and Research Center (MMRRC) at The Jackson Laboratory, an NIH-funded strain repository, and was donated to the MMRRC by David Borchelt, Ph.D., McKnight Brain Institute, University of Florida ^38^. VGAT-Venus mice (B6-Tg(Slc32a1-YFP*)39Yyan) were donated from Dr. Janice Naegele (Wesleyan University, Middletown, CT, USA) ^55^. Mice were housed on a 12 h light/dark cycle, 1-4 mice per cage. Adequate measures were taken to minimize pain and discomfort. The temperature and humidity were controlled, and the cages were individually ventilated. All animal procedures were approved by the Massachusetts General Hospital IACUC (protocol number 2012N000085) and performed under the Public Health Service Policy on Human Care of Laboratory Animals. The study is reported following ARRIVE guidelines.

### Harvesting MGE interneuron progenitors

Donor embryonic medial ganglionic eminence (MGE) interneurons progenitors were obtained as previously described ^60,71^. The four transgenic donor strains including VGAT-Venus, GP5.17, VGAT-ChR2-EYFP, and VGAT-Cre; Ai214 mice were used to harvest MGE cells. Transplantation media consisting of 2 mL Lebovitz’s L-15 media, 20 µL B27, and 1 µL murine EGF was prepared on ice. MGE-IN progenitor was collected from embryonic days (E) 13.5 embryos with the mouse sacrificed in a CO2 chamber. Embryos were placed in sterile ice-cold HBSS -/- and dissected using fine forceps under a dissecting microscope (Zeiss, Discovery. V8). The MGE tissue was then transferred to a 0.6 ml tube containing ice-cold transplantation media and triturated using a P200 pipette to get a cell suspension on ice. The suspension was filtered through a 40 µm filter (Corning, #352340). Dissociated cells were stained with Trypan Blue and counted using a LUNA FL cell counter (Logos biosystem). Dissociated MGE cells were concentrated using a centrifuge for 2 minutes at 800 x g at 4 °C. The cell density was adjusted to the desired concentration (≈500,000 cells/μl) by resuspending the cell pellet in transplantation media.

### MGE interneuron progenitors transplantation

2-month-old (P60, ±7 days) APP mice were anesthetized with isoflurane (5% for induction, 1.5–1.8% for maintenance), and their heads were stabilized in a stereotaxic apparatus. The surgical site was sterilized with 70% ethanol and iodine. Lidocaine (0.1%) was injected subcutaneously at the incision site.

Meloxicam was administrated via the intraperitoneal injection before the surgery. A midline incision was made to expose the skull. The injection sites were determined in the left hemisphere at the following coordinates: AP: +1.4, ML: +1.4, DV: -1.0 mm. A volume of up to 2 µL of cell suspension was injected at a rate of 100 nL/min into burr holes. The Hamilton needle (Hamilton, 26 G, 7804-03 and 80336) was left in place for 5 minutes after injection to allow for the settlement of injected cells. Post-injection, the incision was sutured and mice were allowed to recover on the heat pad. Mice received meloxicam (200 µL) and Tylenol (10 mL) in their drinking water for analgesia for three days following the surgery.

### Whole Brain Imaging with Tissue Clearing

The tissue clearing was performed as previously described ^72,73^. Mice were perfused with ice-cold 50 mL PBS followed by 50 mL 4% PFA. Brain samples were collected and placed in 4% PFA at 4°C overnight for less than 24 hours, then transferred to PBS for another 24 hours at 4°C. The sample was incubated in a hydrogel crosslinking solution consisting of PBS with 4% PFA, 4% acrylamide (Sigma, A3553), 0.02% bis-Acrylamide (RPI, A11270-25.0), and 0.25% VA-044 (TCI, A0312) for 2-3 days at 4°C to allow diffusion of the solution through the tissue. The solution was kept cold before and after adding VA-044 to prevent premature polymerization. After incubation, the sample was placed in a vacuum at 37°C for 3 hours to initiate polymerization using the X-CLARITY polymerization system (Logos Biosystems). The sample was wiped using a paper towel to remove excess hydrogel solution. The sample was then rinsed with 50 mL PBS five times over 24 hours. The sample was delipidated using an active electrophoretic tissue clearing (ETC) system at 37 °C for 24 hours. The clearing solution was circulated through it using a temperature-controlled water circulator. The samples were incubated in a refractive index (RI) matching solution (Easy Index, EI-500-1.52, RI = 1.52) for 24 hours at room temperature with gentle shaking followed by immersion in the fresh solution for another 24 hours. Fluorescence images were collected using a Zeiss Lightsheet Z7 microscope. Image data was reconstructed and visualized using Arivis software (Zeiss).

### Free-Floating Immunohistochemistry (IHC)

Mice were perfused with ice-cold 40 mL PBS followed by 40 mL 4% PFA. Brain samples were collected and placed in 4% PFA at 4°C overnight for less than 24 hours. The sample was immersed in 15% sucrose in PBS for 24 hours followed by 30% sucrose for at least 2 days. Samples were cut 40 μm thick on the coronal plane using a vibratome (Leica). Slice section samples were either used immediately or stored in cryoprotectant at -20°C. Free-floating immunohistochemistry (IHC) was performed based on previously established methods ^74^. Brain sections were transferred into the TBS and rinsed 5 times for 10 minutes each on a shaker to remove the cryoprotectant buffer. The sample was permeabilized and blocked with a blocking buffer consisting of TBS with 3% of normal goat serum (Jackson ImmunoResearch laboratory) and 0.25% Triton-X at room temperature for 2 hours. Tissue sections were incubated with the primary antibody solution at 4°C on a rocking shaker at ∼50 rpm overnight. The following primary antibodies were used at the dilutions with blocking solution including chicken anti-GFP (1: 500; Aves, GFP-200), mouse anti-NeuN (1: 500; Millipore, MAB377), rabbit anti-Iba1 (Fujifilm Wako, 019-19741), rabbit anti-GFAP (1: 1000; Abcam, ab7260), rabbit anti-Olig2 (1: 500; Millipore, AB9610), mouse anti-CaMKII (1: 500; Enzo, ADI-KAM-CA002), mouse anti-GAD67 (1: 1000; Abcam, ab26116), rabbit anti-Ki67 (1: 400; CST, 12202S), rabbit anti-SST (1: 200; Thermo Fisher, PA5-85759), mouse anti-PV (1: 1000; Millipore, P3088), mouse anti-gephyrin (1: 500; Synaptic Systems, 147 011), guinea pig anti-Bassoon (1: 500; Synaptic Systems, 141 318), rabbit anti-VGAT (1: 500; Millipore, AB5062P), mouse anti-PSD95 (1: 500; Millipore, MABN68), host anti-SOX2 (1:500; Abcam, ab97959), host anti-Nkx2.1 (1:500: Abcam, ab76013), host anti-Lhx6 (1:200; Santa Cruz Biotechnology, sc-271433), host anti-Prox1 (1:500; Abcam, ab199359). After washing with 0.25% Triton-X in TBS three times for 10 minutes, sections were incubated with the secondary antibody solution at room temperature on a rocking platform shaker at ∼50 rpm for 2 hours, protected from light. For IHC with primary antibodies derived from the mouse, the sample was blocked from endogenous mouse immunoglobulins with M.O.M.® (Mouse on Mouse) Blocking Reagent (MKB-2213-1, Vector laboratories) at room temperature for 2 hours before being incubated with the primary antibody solution. After washing with 0.25% Triton-X in TBS three times for 10 minutes, sections were incubated with the secondary antibody solution at room temperature on a rocking platform shaker at ∼50 rpm for 2 hours, protected from light. The following primary antibodies were used at the 1: 500 dilutions with blocking solution including (Thermo Fisher, A11004, A11005, A11011, A11012, A11039, A21450, A21235, A31553, and A48255). After washing with TBS three times for 10 minutes, sections were mounted to slide grass using a paintbrush. The tissue was dried using Drierite. After drying, mounting medium (Antifade medium with DAPI (Vectashield, H1500-10) or Prolong diamond RI 1.52 (Thermo Fisher, P36984)) was applied to each slide and covered with glass coverslips and sealed with nail polish. Fluorescence images were collected using a confocal microscope (Olympus, FV3000) or a super-resolution microscope (Zeiss Elyra). SIM images were prepared for the evaluation of synaptic density measured 64 × 64 × 10 μm in size and reconstructed using Zeiss ZEN software. Continuously rendered process-like structures were selected for assessment. The total synaptic lengths analyzed ranged from 10 to 50 μm per sample, with manual observation of 1 to 11 synaptic boutons in each case. Imaging data were analyzed with software including imageJ or Arivis software (Zeiss).

### In Vivo Multiphoton Calcium Imaging

Calcium imaging was performed as previously described. Mice were initially anesthetized with 5% isoflurane and maintained on 1.5% isoflurane during surgery. The mice were placed on a heating pad to maintain body temperature at approximately 37 °C. Ophthalmic ointment was applied to protect their eyes. The skin was removed to expose the skull, and the skull was scrubbed with cotton swabs to remove the membrane. Cranial windows were placed in the anterior cortex and injected with MGE-IN progenitors. A circular hole was created using a surgical drill and drill bit. The dura matter was kept intact and wetted with ice-cold PBS. 5mm windows were mounted and sealed around the outside with a mixture of super glue and dental cement. Meloxicam (5 mg/kg) and acetaminophen (300 mg/100 mL) were administered as post-operative analgesics for 3 days. Two-photon imaging was conducted using a Fluoview FV1000MPE multiphoton microscope (Olympus) with a mode-locked MaiTai Ti sapphire laser (Spectra-Physics). Imaging was performed at least 3 weeks after installation of the cranial window, when the mice recovered and the cranial window condition improved, and at least 2 months after transplantation, when the transplanted cells matured. Mice were sedated with 5% isoflurane in room air using the SomnoSuite® Low-Flow Anesthesia System (Kent Scientific). Imaging was conducted under light anesthesia and low airflow rates (1% isoflurane and ∼40 mL/min airflow for a 30 g mouse). A heating pad maintained the body temperature at 37.5°C. The Fluoview software was controlled for scanning and image acquisition. Spontaneous calcium transients were collected within the somatosensory cortex at 5-10 Hz through a 25x 1.05 numerical aperture water immersion objective (Olympus) at 1-5x digital zoom. Multiple fields of view (approximately 160 × 100 µm, 1 pixel per µm) were imaged per mouse, with each field of view recorded for at least 100 seconds. We used an established MATLAB program (https://github.com/moustaam0/Algamal2022_analysis_w_OASIS) to analyze calcium images calculating event rate ^24^.

### Voltage-Sensitive Dye (VSD) Imaging

VSD imaging was performed as previously described. Imaging was performed at least 3 weeks after the installation of the cranial window after the mice recovered and the cranial window condition improved, and at least 2 months after transplantation, when the transplanted cells matured. Mice were initially anesthetized with 5% isoflurane and maintained on 1.5% isoflurane during surgery. The mice were placed on a heating pad to maintain body temperature. Ophthalmic ointment was applied to protect their eyes. The skin was removed to expose the skull, and the skull was scrubbed with cotton swabs to remove the membrane. Cranial windows were placed over the right somatosensory cortex. A circular craniotomy was created using a surgical drill and drill bit. The dura mater was removed. RH2080 was topically applied to the cortex and incubated for 90 minutes using surgical sponges. Silicon grease was applied to the edges of the craniotomy to avoid leakage during incubation. After incubation, VSD dye was washed off with surgical sponges soaked in PBS. Clean 5mm windows were prepared with isopropyl alcohol and dried. 5mm windows were mounted over the craniotomy and sealed with a mixture of super glue and dental cement. A light-guide cannula (Doric Lenses) was installed above the anterior left cortex over the site of cell transplantation. C&B Metabond (Parkell) was applied to cement at the edges of the surgical area, thereby securing the cannula and cranial window. Voltage-sensitive dye (VSD) imaging was conducted using a CMOS-based fluorescence microscope (Olympus, BX50WI). Optogenetic stimulation was performed during VSD imaging under three distinct illumination conditions. For pulsed-wave illumination at 0.6 Hz, TTL sequences were generated using a DAQ USB device (USB-6001, National Instruments), producing pulses with a duration of 400 ms. Random-wave illumination was controlled by a Raspberry Pi 4 TTL controller (Raspberry Pi), delivering light with a 24% duty cycle and 50 ms timing precision. Custom software developed with the *pittl-client* library was used to operate the TTL controller. Continuous-wave illumination was applied without a TTL controller, delivering continuous-wave light for up to 2 minutes, with a 10-minute recovery period for prolonged applications. Non-illuminated conditions served as the control group. The light source, excitation filter (Chroma, ET630/30m), fluorescence filter (Chroma, ET665LP, and ZET473NF), 2x objective lens, and CMOS camera (Hamamatsu, C13440) were configured to detect fluorescence signal. The microscope was operated using dedicated software (HC Image Live). Imaging was performed with a binning of 2 and a resolution of 256 × 256 pixels. The exposure time was set to 5 ms, capturing 5000 consecutive frames per session. Imaging was conducted either during optogenetic stimulation or under non-illuminated conditions as a control. Data were saved in CXD format for subsequent analysis. We used an already established MATLAB program to analyze VSD images calculating the slow oscillation power ^18,20^.

### Statistical Information

All statistical analyses were performed using GraphPad Prism 10.4.1. Data are presented as the mean ± SEM. For each statistical comparison, normality was assessed with the Shapiro–Wilk test and either parametric or nonparametric tests were chosen accordingly. Nonparametric tests used were either the Mann-Whitney *U* Test or the Kruskal–Wallis test followed by Dunn’s multiple comparisons test depending on the number of groups. When comparing two groups with the parametric test, if the *F*-test indicated that the variances were equal, the Student’s *t*-test was used. If not, the Welch’s *t*-test was used. For comparisons among more than two groups with the parametric test, if the *F*-test indicated that the variances were equal, the one-way ANOVA followed by Tukey’s multiple comparisons test was used. If not, Welch’s ANOVA followed by Dunnett’s T3 multiple comparisons test was used. To examine the effects of two categorical independent variables on a continuous dependent variable, two-way ANOVA was performed and followed by Tukey’s or Šídák’s multiple comparisons test. When more than two conditions are measured repeatedly on the same subjects (e.g., VSD data measured present and absent of optogenetic stimulation in the same mouse), a repeated-measures analysis is used to account for within-subject variability.

## Acknowledgment

This work was supported by the Overseas research fellowship, Japan Society for the Promotion of Science 202360054; Research Fellowship, Japan Agency for Medical Research and Development (AMED); National Institutes of Health Grant R01AG066171 and R21AG075807. We thank Drs. Qiuchen Zhao, Maria Virtudes Sanchez Mico, Steven S. Hou, Masato Maesako, Brian Bacskai, and Stephen N. Gomperts for technical support and advice. We thank the Harvard Center for Biological Imaging (RRID: SCR_018673) for infrastructure and support. Illustrations were created with BioRender.com.

## Author contributions

Conception and design of study (S.Y., M.M., K.V.K.), animal experiments (S.Y., M.M., M.A.), histological experiments (S.Y., A.M.S., M.R.M., S.J.P., S.T., H.B., L.L., A.L., R.L.G., T.J.Z.,), wrote the manuscript (S.Y., D.V., K.V.K.), edited the manuscript (S.Y., D.V., K.V.K.), and project supervision (D.R., J.R.N., D.V., K.V.K.), All authors read and approved the final manuscript.

