## Supplemental Figures for "Transplantation of GABAergic Interneuron Progenitors Restores Cortical Circuit Function in an Alzheimer’s Disease Mouse Model"

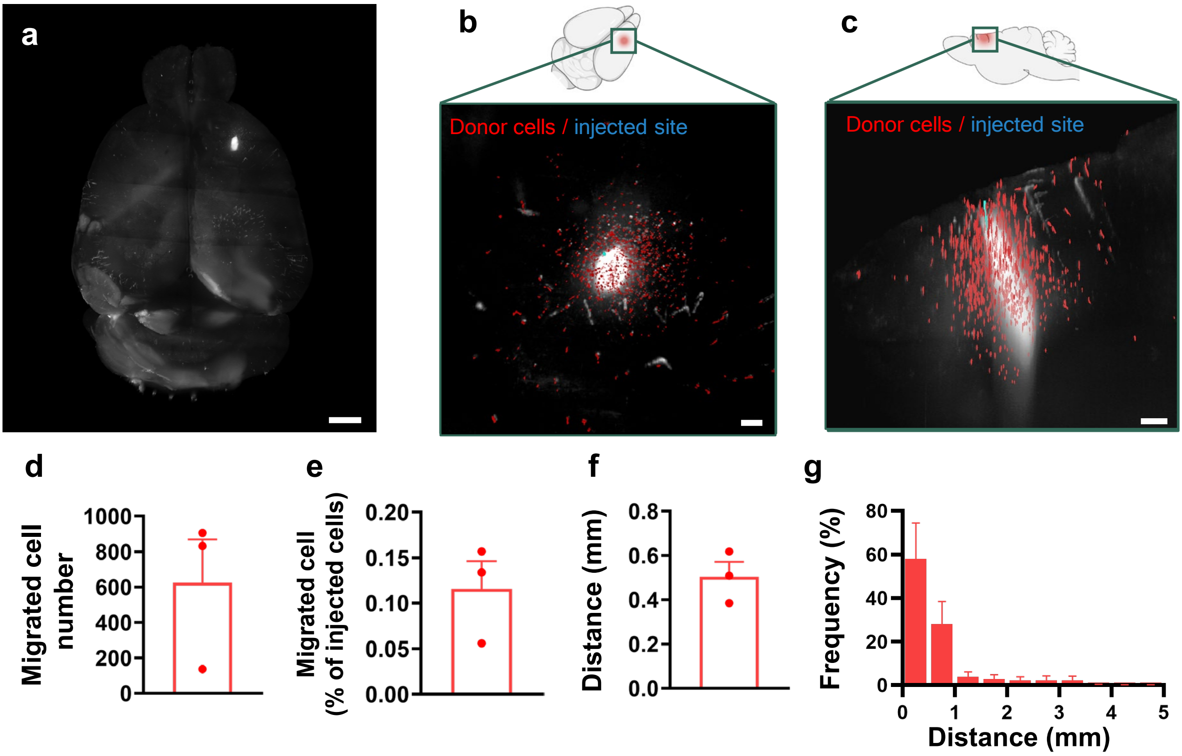


**Supplementary Fig. 1 | MGE cells transplanted into wild-type cortex survive for 60 days and migrate.** **a** Three-dimensional (3D) reconstruction of the entire host brain. **b, c** Higher magnification 3D reconstructions from the dorsal (**b**) and sagittal (**c**) views showing donor cells (red) and the injection site (blue). **d** Number of migrated cells detected by whole-brain imaging. **e** Percentage of donor cells that migrated, calculated by dividing the number of migrated cells by the total number of transplanted cells. **f** Mean distance traveled by donor cells, measured from the injection site. **g** Distribution of distances traveled by donor cells. Data are presented as mean ± SD. Scale bars: 1 mm (**a**), 0.2 mm (**b**), 0.3 mm (**c**). $n = 3$ mice.

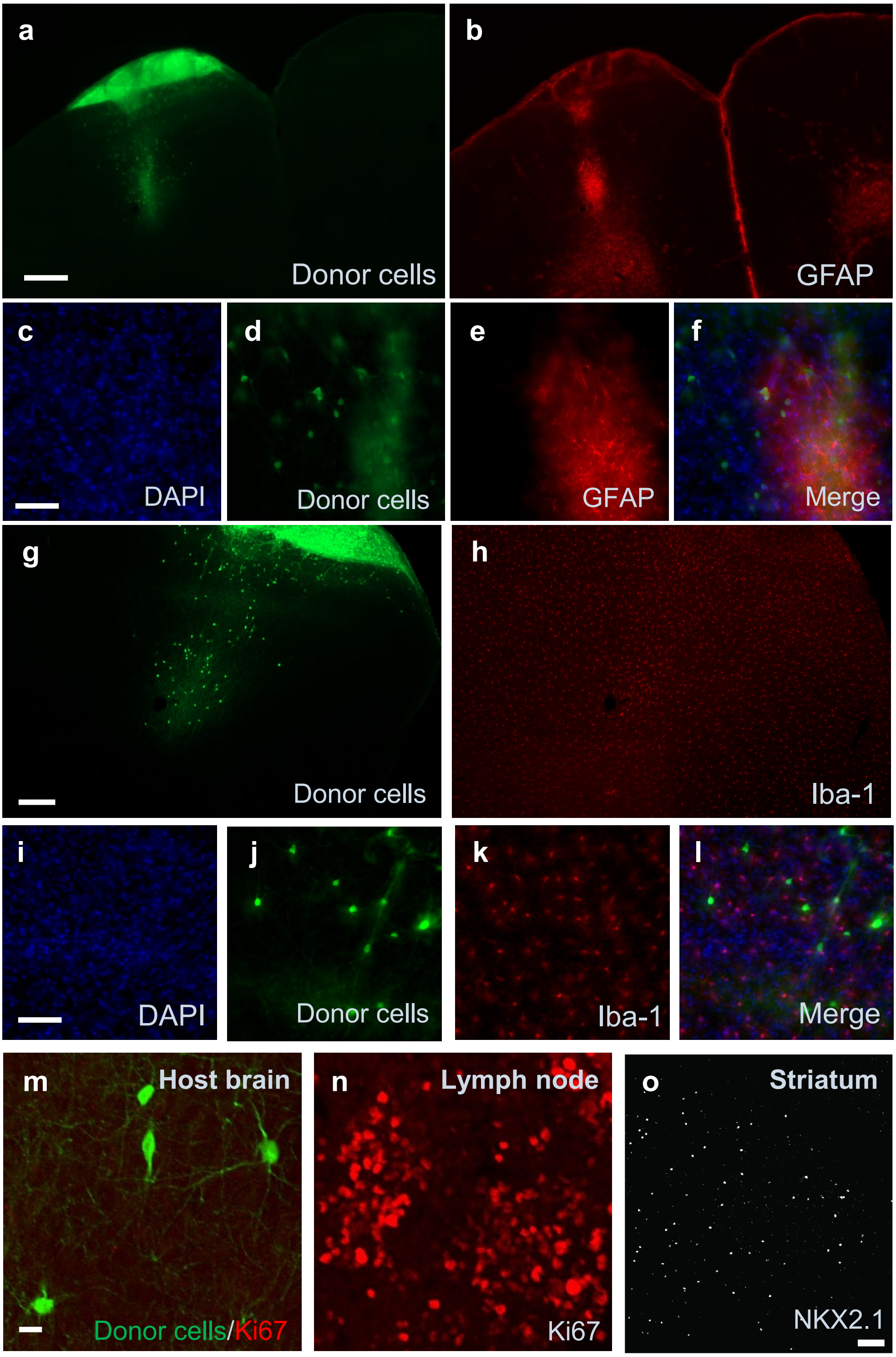


**Supplementary Fig. 2 | Immunostaining revealed GFAP-positive astrocyte accumulation, but not microglial accumulation, at the injection site a–f** GFAP immunostaining in the host anterior neocortex 2 months post-transplantation. **g–l** Iba-1 immunostaining in the host anterior neocortex 2 months post-transplantation. **m** Ki67 immunostaining in the host anterior neocortex 2 months post-transplantation. **n–o** Staining quality controls: lymph node for Ki67 (**n**) and striatum for NKX2.1 (**o**). Scale bars: 400 µm (**a**), 50 µm (**c, i**), 20 µm (**m**), and 100 µm (**o**).

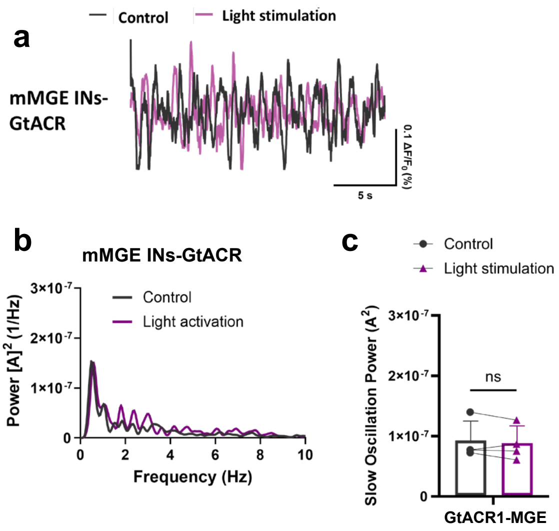


**Supplementary Fig. 3 | Random light stimulation of transplanted MGE cells does not affect slow oscillation power. a** Raw fluorescence traces from the host APP/PS1 cortex during random light stimulation (purple) and no stimulation (gray). **b** Power spectral density of MGE-transplanted APP/PS1 cortex with or without optogenetic stimulation $[A]^2$ = magnitude of Fourier amplitude squared. **c** Slow oscillation power with or without optogenetic light stimulation. Each data point represents the average of 10–15 traces from each mouse. Data are presented as mean ± SD, with ns indicating no significant difference. $n = 4$ mice.
